# Retinal Resistivity Mapped by Electrical Resistance Tomography

**DOI:** 10.64898/2026.10.13.756866

**Authors:** Anna Kochnev Goldstein, Mohammad Asif Zaman, Michael Aaron Sommeling, Sarthak Vinay Shah, Amrith Lotlikar, Mohajeet B. Bhuckory, Davis Pham-Howard, Zhijie Charles Chen, Praful Vasireddy, A.J. Phillips, Aviv Sharon, E.J. Chichilnisky, Daniel Palanker

## Abstract

Retinal resistivity is a key parameter for modeling both the electric field in tissue and the resulting neural response - a crucial step in designing retinal prostheses, especially since both depend on resistivity nonlinearly. Previous studies using penetrating microelectrodes reported vastly different values, varying by as much as two orders of magnitude, likely due to parasitic capacitance in microelectrodes, compression of tissue, and its rupture by the inserted probe, allowing the conductive medium to seep in and alter the measurement. To measure retinal resistivity without compression or rupture, we applied electrical resistance tomography using a planar multielectrode array with a 30 µm pitch. Voltage waveforms generated in response to current injected from one electrode were measured on other electrodes in the array, at distances ranging from 30 to 600 µm. Data from wild-type and degenerated rat retinas were analyzed using two methods: a commercial inversion software, Res2DInv, and a custom machine learning-based decoder, with healthy retina approximated as a 4-layer structure and degenerated retina - as a 3-layer one.

In healthy retina, average resistivity was 2.7 kΩ·cm for the combined ganglion cell-nerve fiber layer, 1.6 kΩ·cm for the inner plexiform layer, 2.3 kΩ·cm for the outer and inner nuclear layers, and 1.0 kΩ·cm for the photoreceptor inner and outer segments; a single-layer approximation yielded 1.7 kΩ·cm. In degenerated retina, average resistivity was 3.4 kΩ·cm for the ganglion cell-nerve fiber layer, 1.1 kΩ·cm for the inner plexiform layer, and 4 kΩ·cm for the inner nuclear layer, with a single-layer approximation of 2 kΩ·cm.

**Significance:** Designing retinal prostheses to restore vision after loss of photoreceptors requires accurate modeling of electric field in tissue, but published resistivity values vary by two orders of magnitude due to inconsistencies with penetrating microelectrode-based measurements. We adapted electrical resistance tomography, a non-destructive technique from geophysics based on inverse problem solving, and developed a machine learning-based decoder to map resistivity in rat retinas using a planar microelectrode array. Our analysis yields quantitative, layer-specific resistivities for healthy and degenerated retina. Beyond retinal prostheses, this approach offers a general, deformation-free strategy for measuring electrical properties of biological tissues.

---

Retinitis Pigmentosa (RP) and Age-Related Macular Degeneration (AMD) are among the leading causes of untreatable visual impairment. However, despite the irreversible loss of photoreceptors in retinal degeneration, the inner retinal neurons survive to a large extent [1], [2], [3], providing an opportunity to partially restore sight by electrical stimulation with a retinal prosthesis [4]. Design of such prostheses heavily relies on modeling the neural stimulation with various electrode configurations and pulsed waveforms, which assumes certain electrical properties of the tissue and the electrodes. One of the most important parameters to be defined is the electrical resistivity of the retina. Unfortunately, the values reported in previous studies vary by nearly two orders of magnitude, ranging from approximately 59 to 8000 Ω·cm (Table S1, *Supplementary Materials*), making reliable quantitative modeling very difficult. Moreover, while some studies only report one resistivity without addressing the anatomical differences between the retinal layers [5], others map the resistivity as a function of depth without relating the obtained values to specific layers, which limits their practical utility [6], [7], [8].

Most previous studies relied on penetrating microelectrodes advancing through the tissue to measure impedance at various depths [6], [7], [8], [9]. Since these electrodes puncture the retina, they allow the highly conductive perfusion media to reach the formed cleft, reducing the local resistivity. At the same time, the penetrating electrode compresses the tissue in front of it, resulting in reduced water content and increased resistivity due to denser packing of cells [10]. Some studies based on Electrical Impedance Spectroscopy reported impedance measured at only one frequency, without providing any data on the rest of the spectrum, which is essential to parse out the resistance and capacitance at the electrode-electrolyte interface [5], [11]. Other studies did consider the spectrum, but extracted the access resistance at a frequency where the phase angle was still around 20 [7] or even 55 degrees [6] rather than 0, indicating a significant contribution of non-resistive components in the assumed Randles circuit. Since access resistance is the only component in the Randles circuit indicative of the retinal resistivity, extracting a reliable value that is not contaminated by capacitance is a general problem with small electrodes.

To overcome these limitations, we adopted an approach that does not physically alter the tissue’s morphology - Electrical Resistance Tomography (ERT) using a multielectrode array (MEA) with planar (non-penetrating) probes of a 30 µm pitch and no tissue compression. ERT is commonly used in geophysics to characterize the resistivity of the earth’s layers and has also shown potential in several medical applications, such as monitoring pulmonary ventilation and gastric emptying [12], though only qualitatively. Here, we applied the same technique, albeit scaled down in size by about a million, to micrometer-scale dimensions of the retina.

As modeling for retinal prostheses usually focuses on degenerated retina, we performed ERT on samples not only from healthy Long Evans (LE) rats but also the Royal College of Surgeons (RCS) rats with degenerated retina. The resistivity profile of the measured samples was decoded using two different methods:

1) Res2DInv, a commercial inversion software widely used in geophysics, and 2) a custom Machine Learning (ML)-based decoder with training data generated from a COMSOL Multi-physics model. The former is a non-parametric method, while the latter requires an assumption of the number and thickness of distinct layers, which we obtained from retinal imaging. We quantify the resistivity of 4 retinal layers in healthy retina, 3 layers in the degenerated retina, as well as the average resistivity for the whole retina in both cases, and discuss the advantages and limitations of each decoding method.

## Methods

### Animals and Tissue Preparation

All experimental procedures were conducted with retinas harvested from terminally anesthetized rats in accordance with the Statement for the Use of Animals in Ophthalmic and Vision research of the Association for Research in Vision and Ophthalmology (ARVO) and approved by the Stanford Administrative Panel on Laboratory Animal Care (APLAC). Animals were maintained at the Stanford Animal Facility under 12 h light/12 h dark cycles with food and water ad libitum.

Two types of retinas were studied: (1) healthy Long Evans (LE) rats and (2) Royal College of Surgeons (RCS) rats after a complete degeneration of photoreceptors. Following enucleation, the eyes were placed in an oxygenated Ames solution (Sigma Aldrich) at 33°C, hemisected, vitrectomized, and pinned for stability. Retinal pieces of 3-4mm in diameter were created and detached from the pigment epithelium. Overall, 9 retinal samples from 9 LE rats (7 from the right eye and 2 from the left; 6 male, 3 female; average age 256 ± 77 days; 3 ventral, 2 dorsal, and 4 undetermined) and 6 samples from 6 RCS rats (3 from the right eye and 3 from the left; 4 male, 2 female; average age 359 ± 90 days; 4 dorsal, 1 ventral, and 1 in-between) were used in these studies. Each measurement was completed within two hours after enucleation.

### Retinal Layers

Histology and optical coherence tomography (OCT) delineated the following layers in healthy LE rats (Figure 1A): Inner and Outer Segments of photoreceptors (IS + OS), Outer Nuclear Layer (ONL), Outer Plexiform Layer (OPL), Inner Nuclear Layer (INL), Inner Plexiform Layer (IPL), Ganglion Cell Layer (GCL), and the Nerve Fiber Layer (NFL). In RCS rats, the photoreceptors are missing, so only INL, IPL, GCL, and NFL are visible (Figure 1B). OCT can provide the thickness of the different layers in-vivo, and we used those as priors for decoding the retinal resistivity. The overall average thickness of a healthy retina and a degenerated retina was measured to be about 200µm and 126µm, respectively.

**Figure 1:**
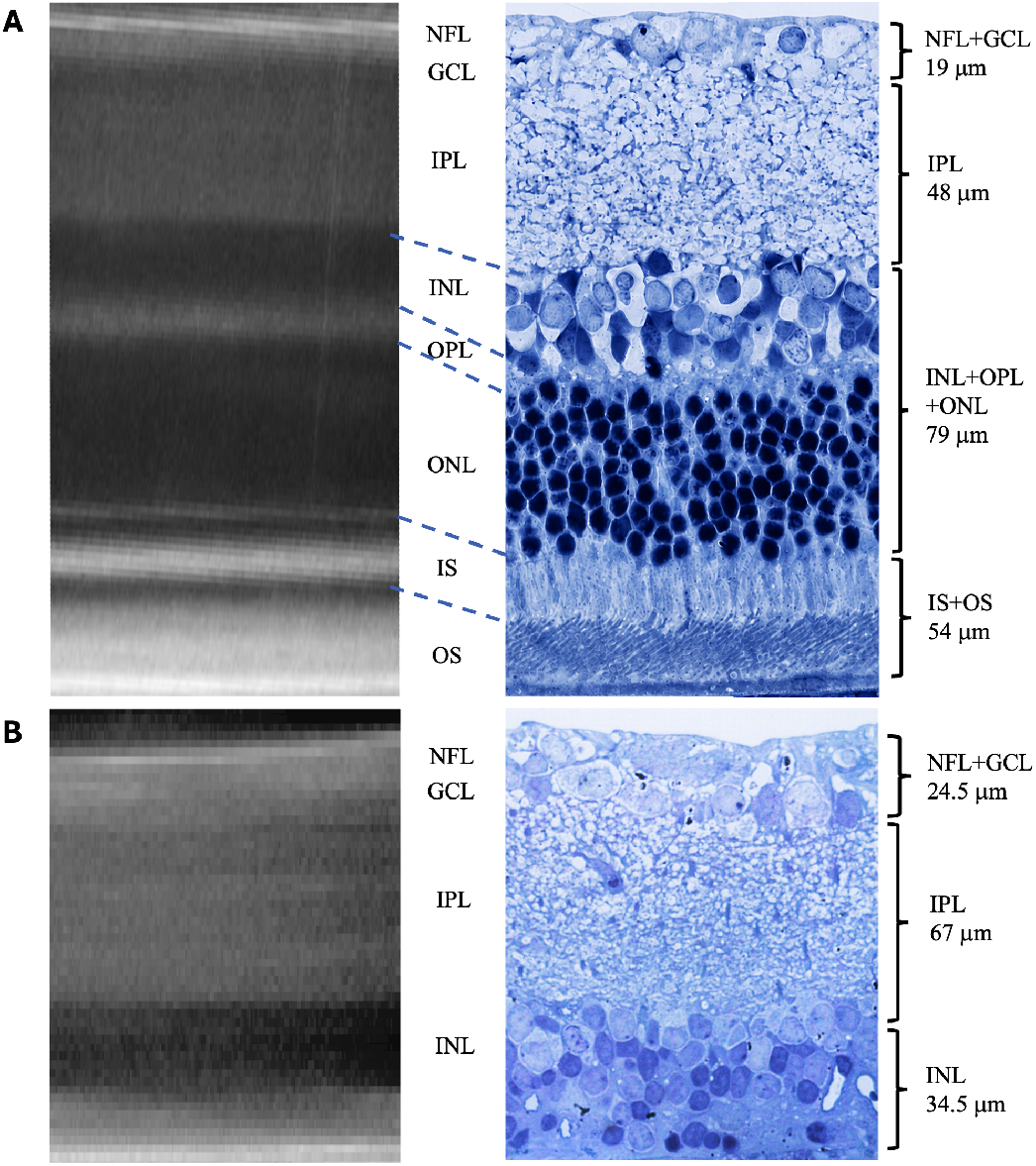
Retinal layers in OCT and histology. **A)** OCT (left) and histology (right) of a healthy LE rat retina (images obtained from two different animals). For decoding purposes, the layers are combined into four groups: NFL+GCL, IPL, INL+OPL+ONL, and IS+OS. **B)** OCT obtained at the age of 295 days (left) and histology (right) of a degenerated RCS rat retina. For decoding purposes, the layers are combined into three groups: NFL+GCL, IPL, and INL. The layer thicknesses are a population average and not representative of the given animal.

The depth resolution in ERT is limited by the minimal pitch of the measurement system [13]. Since the OPL, GCL, and NFL are thinner than the minimal pitch of our MEA (30 µm), they cannot be resolved with our setup and will be masked by the surrounding layers. Therefore, we grouped these layers with their neighbors: OPL was grouped with the nuclear layers – INL and ONL, and GCL was grouped with NFL. This resulted in a 4-layer model for the LE rat and a 3-layer model for the RCS rat (Figure 1). Since many practical modeling approximations would benefit from a single resistivity value, we also analyzed a 1-layer model. It is important to note that even after grouping the GCL with the NFL, the combined thickness remains below 30 µm. However, the decoding of the experimental data using the Res2DInv software showed a clear difference between this layer and the subsequent IPL, justifying a 4-layer model for LE and a 3-layer model for RCS despite this limitation.

### Data Acquisition

A hexagonal multielectrode array of 800 µm in width with 512 planar electrodes was used for the measurements [14], [15]. The electrodes were 5 µm in diameter spaced by 30 µm along the vertical axis and by 33.54 µm along the diagonals (Figure 2A). Each experiment began with electroplating platinum black on the electrodes, which would increase their diameter to approximately 10-15 µm. The return electrode was a ring-shaped platinum wire around the circumference of the measurement chamber (Figure 2B).

**Figure 2:**
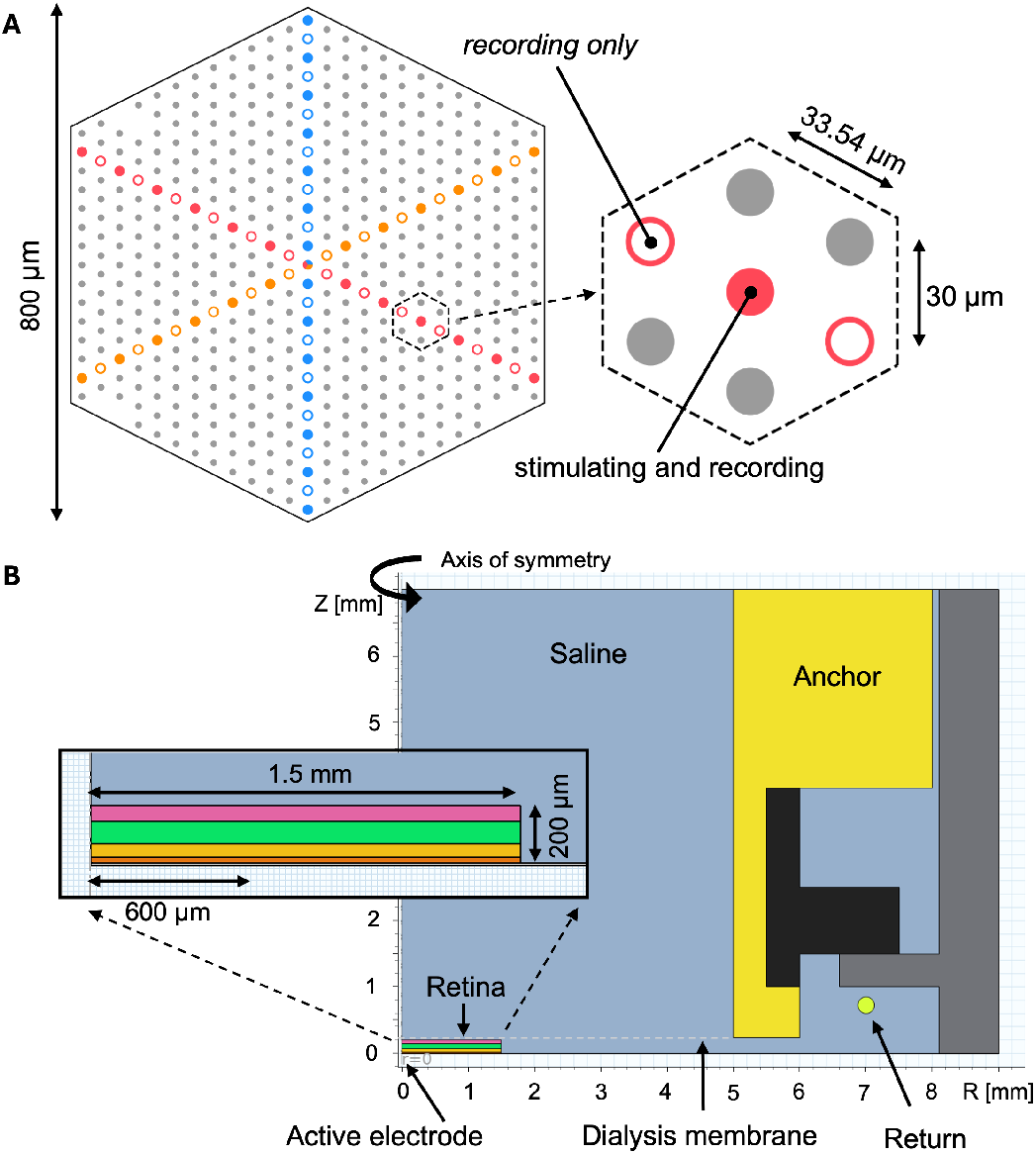
Experimental setup. **A)** Diagram of the MEA – 800 µm in width, with a vertical pitch of 30 µm and a diagonal pitch of 33.54 µm. The three lines used for the measurements are highlighted. Solid circles indicate electrodes used for both stimulation and recording. Hollow circles indicate electrodes used for recording only. **B)** Side view of the axially symmetric COMSOL model of the experimental setup. Insert illustrates a close-up of the 4-layer LE retina, surrounded by PBS, with the injecting electrode on axis of symmetry.

Dissected retinal punches were positioned on top of the MEA. Six LE samples and six RCS samples were measured with the GCL facing down, toward the electrodes, and three LE samples were measured with the photoreceptor layer facing down. These will be referred to as GCL-down and PR-down, respectively. A dialysis membrane stretched over a custom anchor was lowered on top of the tissue to hold the sample in place (Figure 2B). To avoid compressing the sample, the micromanipulator controlling the height of the membrane was calibrated to the average natural thickness of the tissue. Each sample was observed through an IR microscope during anchoring to confirm contact with the membrane and flattening over the array.

In GCL-down samples, after anchoring the retina in place, spiking was briefly recorded to confirm ganglion cell activity. Then, the sample was disconnected from the oxygenated culture medium supply and replaced with Phosphate Buffered Saline (PBS) until the spontaneous spiking rate decayed to zero and displayed no light response (around 10 min) to ensure ERT was not contaminated by spontaneous firing of the retinal ganglion cells. The ERT scan was initiated immediately after the spiking activity subsided.

To retrieve information as far from the current injecting electrode as possible while also providing the highest possible depth resolution, measurements were conducted along the longest and densest three lines (where the electrode pitch was 30 µm and 33.54 µm), as illustrated in Figure 2A. Within each line, one electrode at a time was used to inject a square bi-phasic pulse of current, while other electrodes along that line recorded the voltage. All electrodes outside that line were turned off to minimize parasitic pathways to the ground. The injected current was swept from 0.5-58 nA and consisted of a 0.5 ms anodic phase followed by a 0.5 ms cathodic phase. After applying one current level, the injection site was shifted to the next electrode. After iterating over all the injection sites on the measurement line, the current was increased and the procedure was repeated. Twenty trials were performed for each electrode-current pair.

The pulse duration was chosen to balance between (a) ensuring that the voltage was measured at a sufficiently high frequency, where the impedance is dominated by the access resistance rather than the capacitance, and (b) providing sufficient time for current redistribution over the array to reach a steady state before the pulse is switched off (Figure 3). Although biphasic 0.5ms-long pulses mostly satisfied these conditions, some recordings at long distances (> 300 µm) and in a very resistive calibration medium (> 3000 Ω·cm) did not reach the steady state by the end of 0.5 ms. At higher current levels, the waveforms drop slightly after reaching the peak due to shunting of current at the injection electrode, which was calibrated as described in *Supplementary Materials*.

**Figure 3:**
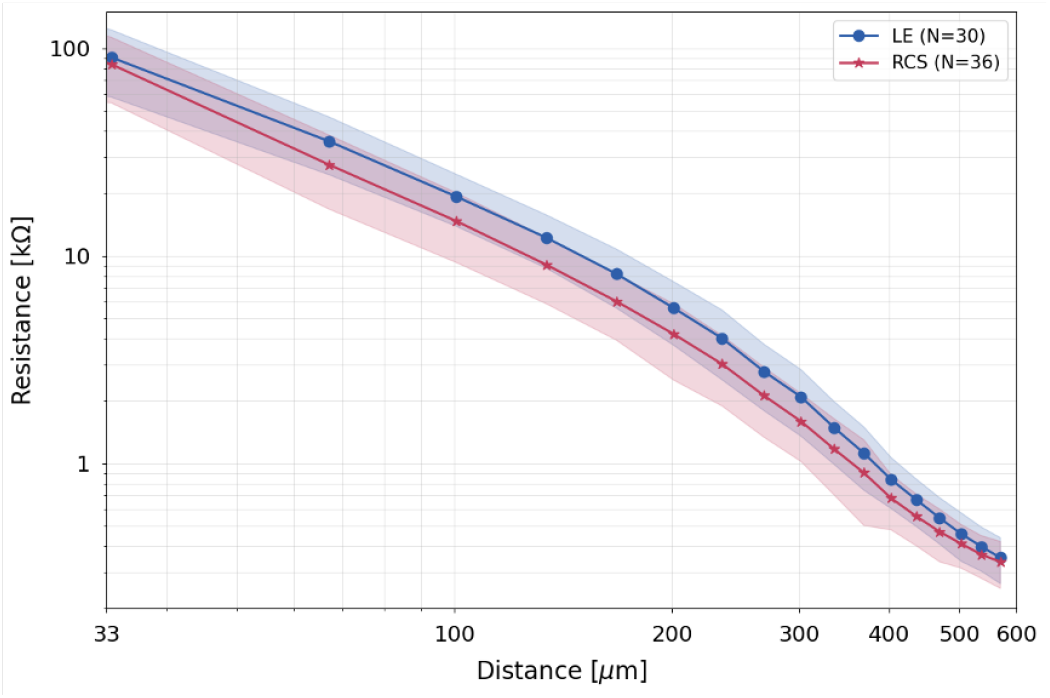
Average resistance-distance traces. Blue: Average LE GCL-down trace (6 samples, 30 traces). Magenta: Average RCS GCL-down trace (6 samples, 36 traces). The shaded band marks ± std from the average.

### Data Calibration and Processing

The peak voltage during the anodic phase was extracted from the raw waveforms for every current level and every pair of injecting and recording electrodes. To average over multiple current levels, a ratio of the measured voltage to applied current was calculated, representing the resistance at a given distance from the injecting electrode. As the recording electrodes were located at different distances, each injecting electrode yielded a series of resistance-distance data points. Resistance values recorded at identical distances were averaged.

To calibrate the measurement system, data was first acquired in a series of diluted saline solutions of known resistivities. To check for possible flaking of brittle Pt black from electrodes during mounting of the retina, calibration was performed before and after the retinal measurement. The calibration procedure, described in *Supplementary Materials*, adjusted for current shunted at the injection electrodes and for gain variability between the amplifiers of the recording electrodes. Three measurement lines on the MEA and two calibrations performed on each line yielded up to six traces per sample. The resulting resistance values across all traces of all samples (of a certain rat type and orientation) were averaged, yielding a single trace which will be referred to as the resistance-distance trace. This trace consisted of 17 data points that could be extracted reliably above the noise level, covering up to 600 µm from the source (Figure 3).

### Resistivity Decoding

To extract the retinal resistivity profile as a function of depth, we applied two decoding methods: a commercial inversion software, Res2DInv (Seequent, The Bentley Subsurface Company, Christchurch, New Zealand, version 2026.2.1.1655), and a custom ML model, which will be referred to as the ML decoder. In both cases, the input data was the average calibrated resistance-distance trace (Figure 3). Res2DInv was not given any prior information about the structural profile and was used as a non-parametric model, while the ML decoder was trained on a fixed number of layers and thicknesses. Due to low sensitivity of the measurement to inner retinal layers when a more conductive layer of photoreceptors was facing the MEA, only the GCL-down orientation could be decoded reliably, as described in details in *Supplementary Materials (Layers Sensitivity Analysis)*.

#### Res2DInv

Res2DInv is a commercial inversion software commonly used by geophysicists to convert voltages measured with a string of electrodes on the surface of earth to a 2D map of the underlying resistivity profile. The input is a list of measurement coordinates (different combinations of 3 coordinates: one injection site and two recording sites) and the corresponding resistances obtained at those locations. To accommodate this format, the 1D experimental resistance-distance trace was transformed into a 2D input by replicating the resistance values for all injecting electrodes.

The output of the software was a 2D map of the decoded resistivity as a function of lateral distance and depth. A 1D resistivity profile was then extracted from the output by horizontal averaging of the resistivity values around the center of the map (from 300 to 500 µm) at each depth, yielding the curve shown in Figure 4B. To provide a numerical estimate for the distinct retinal layers, a single value was extracted within the boundaries of each layer using the peak value or the average over the layer (more details in *Supplementary Materials*). The boundaries used for this numerical analysis matched the 4-layer model for LE rats and the 3-layer model for RCS rats.

**Figure 4:**
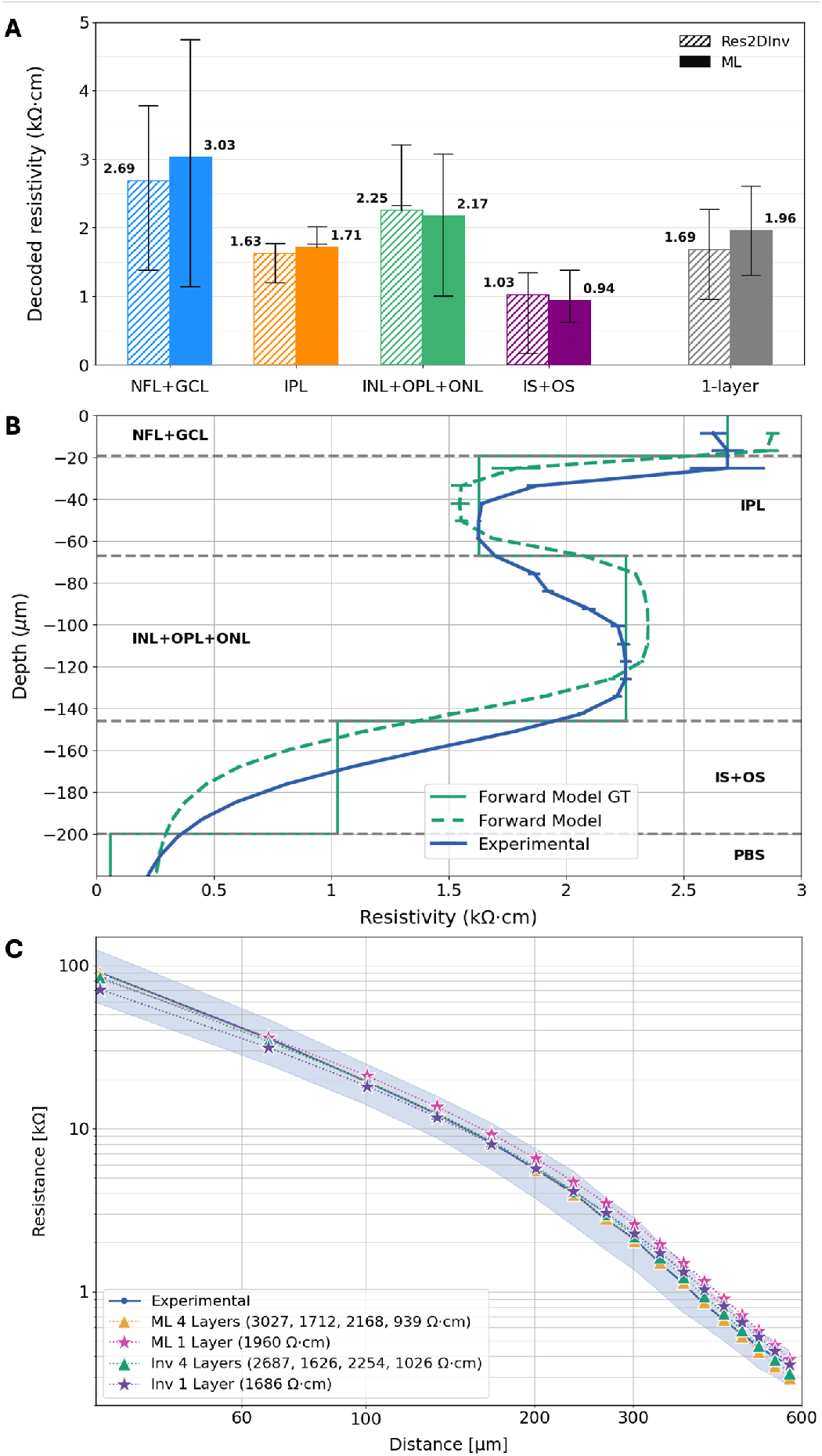
Resistivity of healthy LE retina. **A)** The columns indicate the values for each layer decoded by two methods from the average experimental GCL-down trace. The vertical bars represent the values decoded for the minus and plus std traces. **B)** Res2DInv’s 1D output for the experimental trace, along with the output for the COMSOL forward-model trace generated based on the extracted values: 2687, 1626, 2254, 1026 Ω·cm, and PBS of 60 Ω ·cm, shown by the ground truth (GT) trace. **C)** Experimental average resistance-distance trace, with a shaded band marking the standard deviation, overlaid with the COMSOL forward-model traces generated based on the values extracted with both methods.

#### ML Decoder

The retinal resistivity profile was also deciphered using a custom ML decoder consisting of a 1D Convolutional Neural Network (CNN) followed by three fully connected feed forward layers (Multi-Layer Perceptron). It was trained to take a vector of 17 resistance values sorted by their distance from the injecting electrode and output the resistivity values of the assumed retinal layers. Two separate models were trained for each rat type – a 1-layer model and a multi-layer model.

The training data was generated by simulating the electric field in a layered resistive medium surrounded by saline using COMSOL Multiphysics, a commercial finite element solver. Geometry used for these simulations was based on dimensions of the experimental setup and matched the positions of a line of electrodes in the array (Figure 2B). Current was injected from the center electrode and voltages on the surface of the medium were sampled at various distances with increments of 33.54 µm. The voltage measured by the recording electrode divided by the injection current yielded a resistance value for each distance. The training dataset consisted of 30,000 synthetic resistance sequences for the multi-layer model and 12,000 sequences for the 1-layer model. Each training sequence was obtained by simulating a set of randomized layer resistivities from 60 to 6,000 Ω·cm.

#### Validation of the decoded resistivity

To assess whether the specific values obtained for the LE and RCS profiles by either method in fact replicate the observed experimental traces and thus can be considered reliable, COMSOL forward-modeling was employed. The resistivity values extracted for each profile were fed to the COMSOL model, yielding the corresponding resistance-distance curve to be compared with the experimental trace. A broader accuracy analysis of the decoders was performed separately on a test set of 8,000 synthetic traces, as described in *Supplementary Materials*.

## Results

### Experimental Data

Figure 3 shows the average resistance-distance traces across six LE and three RCS samples. The average trace for each rat type was obtained by averaging all traces across all samples, measurements lines, and calibrations (30 and 36 traces for LE and RCS, respectively). The shaded area around each trace indicates plus and minus one standard deviation from the mean. These traces served as the input data for the two decoding algorithms.

### Decoded Retinal Resistivity

#### Healthy LE Retina

Given the average experimental trace of the LE samples (blue solid line in Figure 3), the ML decoder produced four values of retinal resistivity for the 4-layer model, and one value for the 1-layer model. The 4-layer values are: 3027, 1712, 2168, and 939 Ω·cm for NFL+GCL, IPL, INL+OPL+ONL, and IS+OS, respectively (Figure 4A).

While the ML decoder produces discrete values for each layer by design, Res2DInv outputs a semi-continuous curve of resistivity as a function of depth, shown in Figure 4B. As can be seen, the software produces gradual transitions rather than step-wise boundaries between the layers. A resistivity value was then assigned to each layer as previously described. The 4-layer average values are: 2687, 1626, 2254, and 1026 Ω·cm (Figure 4A). A COMSOL forward model based on these values yielded a resistance-distance plot, which was fed back into Res2DInv, and its output is also plotted in Figure 4B. The latter appears to be a slightly downward-shifted version of the former, showcasing the error introduced by discretizing the semi-continuous output of the software into distinct uniform layers.

To understand the spread in resistivities corresponding to individual samples and measurement lines, traces representing plus and minus one standard deviation from the mean (the lower and upper limits of the shaded blue band in Figure 3) were also decoded. The results are shown in Figure 4A as the black vertical bars. The 1-layer equivalent resistivity from Res2DInv’s output was calculated as a thickness-weighted average of the decoded profile.

For both methods, the COMSOL forward models based on the decoded values show a good correspondence with the experimental trace (Figure 4C), even for a single-layer approximation, but the fit further improves with the 4-layer model. This indicates that while using a single value is convenient for many modeling applications, it does not convey the full complexity of the retinal structure.

As can be seen in Figure 4A, resistivity of the nuclear layers (NFL+GCL and INL+OPL+ONL) is higher than that of IPL and IS+OS. This result is somewhat intuitive as the nuclear layers structurally resemble closely packed spherical particles enclosed within very resistive cell membranes. The large spread (621-4741 Ω·cm with the ML decoder and 939-3027 Ω·cm with Res2DInv) might be due to the inherent variability between animals, decoding accuracy, and limited sensitivity of ERT. However, this spread is much narrower than the previously reported range of 59-8000 Ω·cm.

In many applications, a 1-layer approximation of the retina is preferred for practical considerations. The average 1-layer approximation of 1.7-2.0 kΩ·cm decoded here is higher than the commonly assigned values (0.6-1 kΩ·cm [16], [17], [18]), although the lower bound of the decoded spread (0.96 kΩ·cm) does lie in this range.

#### Degenerated RCS Retina

The same decoding procedure was applied to the experimental RCS trace (magenta in Figure 3). The decoded values, shown in Figure 5A, follow the same trend observed in LE retina, where the two nuclear layers are more resistive than the IPL. However, in the RCS retina, the IPL is slightly less resistive than its LE counterpart, likely due to reduced connectivity between bipolar and ganglion cells in a degenerated retina. Despite the larger variation between layers, the average resistivity for the RCS retina remains about 1.6-1.9 kΩ·cm, similar to LE.

**Figure 5:**
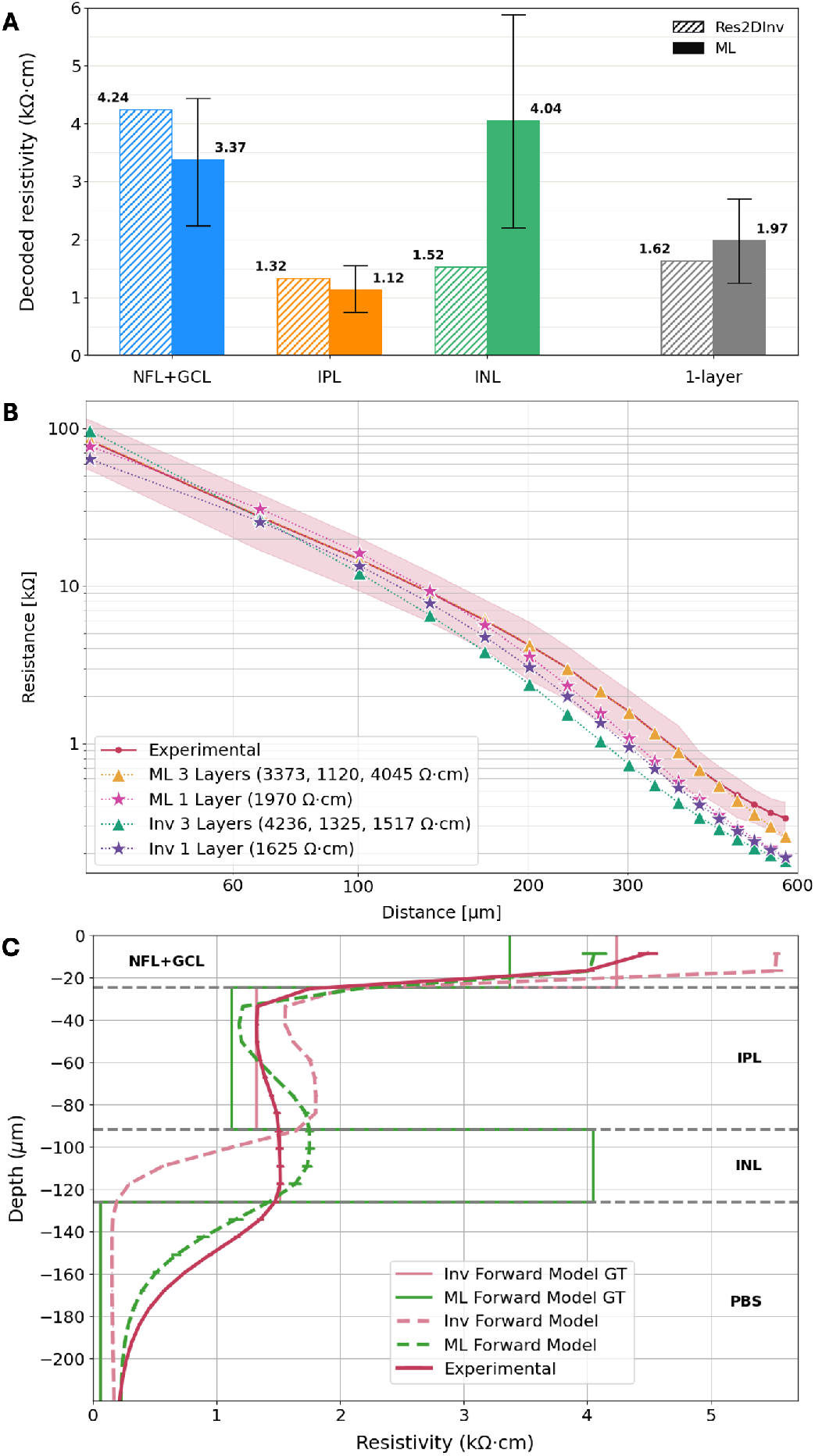
Resistivity of degenerated retina. **A)** The columns indicate the values for each layer decoded from the average experimental GCL-down trace. The vertical bars represent the values decoded for the minus and plus std traces. **B)** Experimental average resistance-distance trace, with a shaded band marking the standard deviation, overlaid with the COMSOL forward-model traces generated based on the values extracted with both methods. **C)** Res2DInv’s inversion of the experimental trace along with its forward-model trace (4236, 1325, and 1517 Ω·cm) and the ML-decoded forward-model trace (3373, 1120, and 4045 Ω·cm). The ground truth input values are shown by the GT traces.

The 3-layer forward-model traces based on the ML-decoded values show good match with the average experimental curve, unlike the 1-layer approximation, which deviates significantly (Figure 5B). The greater deviation might be due to thinner structure and wider variation in the layers thickness between samples and retinal regions, leading to larger animal-to-animal discrepancy. As our model assumes the same layer thicknesses for all animals, it becomes less accurate for RCS compared to LE.

Figure 5B shows that the values decoded by Res2DInv for the 3-layer model do not fit the experimental curve as well. Figure 5C further illustrates how Res2DInv struggles with steep transitions in resistivity between the layers. The output for the forward model based on the extracted values (pink dashed line) rises and drops much faster than the experimental curve (pink solid line). The forward model based on the ML-decoded values follows the experimental curve much closer (dashed green line). This signifies that while Res2DInv by itself cannot resolve a thin and very resistive INL layer between two more conductive layers, it endorses the values suggested by the ML decoder.

## Discussion

Both ERT decoding methods - the commercial Res2DInv software and the custom ML decoder - were in good agreement on resistivity of healthy LE retina and showed that the nuclear layers are more resistive than the IPL and IS+OS layers (2.2-3 vs. 1-1.7 kΩ·cm). The average resistivity of the LE retina as a 1-layer approximation was about 1.7 kΩ·cm. The extremely low (32-59 Ω·cm), saline-like, resistivities previously reported for rats [8] and the very high resistivities (6800-7900 Ω·cm) reported for the frog [9] were not reproduced.

While the forward model for a 1-layer LE retina was a reasonable fit to the measured trace, the deviation further decreased with a 4-layer model. This indicates that while assuming a homogenous retina is convenient for modeling, it fails to convey the full complexity of the multi-layered structure. Moreover, while the 4-layer model yielded a very close fit to the measured trace, even this model did not fully capture the experimental trace.

In RCS retina, the ML decoder yielded a good agreement with the 3-layer model, and the obtained values were significantly higher than the previously reported range of 32-176 Ω·cm [8]. Both methods failed to accurately match the experimental trace with a 1-layer model. This might be due to thinner samples and wider variations in resistivity of different layers in degenerated retina, which makes a 1-layer approximation less accurate. IPL in RCS retina was slightly more conductive than in LE (1.1 vs. 1.7 kΩ·cm) probably due to reduced connectivity.

The utilized decoding methods are complimentary in their advantages and limitations. Res2DInv is a non-parametric method that does not require a model assumption, and it produces a continuous resistivity profile that can offer unbiased insights into the underlying structure. However, it smears sharp transitions and masks steep changes. Therefore, while Res2DInv can be used without an external model, it must be carefully validated on any given profile and when extracting specific numerical values. The ML decoder, on the other hand, assumes a certain model with defined layer thicknesses, and needs to be retrained for any new layered profile or MEA. However, with layer thickness provided by imaging, it produces easily interpretable numerical outputs and performs reliably across steep transitions and variable profiles.

Within the limitations of inverse problem solving, ERT offers a practical approach to measuring the resistivity of biological tissues without causing compression, rupture, and other physical deformations that might affect the result. Moreover, while many resistivity studies did not provide quantification of individual layers and many ERT applications rely on comparison to a known ground-truth or the detection of spatiotemporal changes, this work paves the way for a broader utility that relies on quantitative analysis.

## Data Availability

The software framework used to process the experimental data and run the ML decoder was implemented in Python and is publicly available at: 10.5281/zenodo.23351971, along with the COMSOL model and the settings file used for the Res2DInv inversion.

## Author Contributions

A.K.G, S.V.S, M.A.Z, M.A.S, A.L, Z.C.C, and D.P developed the methodology. D.P and E.J.C provided the resources. M.A.S, A.L, P.V, A.J.P, and A.S conducted experiments and collected data. A.K.G developed the data processing pipeline and processed the data in Res2DInv. M.A.Z developed the COMSOL model and the ML decoder. M.A.Z and D.P.H performed OCT imaging. M.B analyzed OCT and histological images. A.K.G and S.V.S conducted the literature review. A.K.G wrote the manuscript. A.K.G, M.A.Z, M.A.S, and D.P.H prepared the figures. All authors reviewed and edited the manuscript.

## Acknowledgments

We thank Paweł Hottowy, Alexander Sher, and Alan Litke for developing the 512-electrode system and for related technical assistance; Leonid Ryzhik for insightful discussions of the inverse problems; Subhasish Mitra for general support, and the Seequent Res2DInv support team for technical assistance and useful discussions. Studies were supported by the National Institutes of Health (Grants R01-EY-035227, and P30-EY-026877), the Department of Defense (Grant W81XWH-22-1-0933), AFOSR (Grant FA9550-19-1-0402), and an unrestricted grant from the Research to Prevent Blindness.

## References

[1] J. L. Stone. Morphometric Analysis of Macular Photoreceptors and Ganglion Cells in Retinas With Retinitis Pigmentosa. Archives of Ophthalmology, 110(11):1634, November 1992.

[2] M. S. Humayun, M. Prince, E. de Juan, M. Moskowitz, I. B. Klock, and A. H. Milam. Morphometric Analysis of the Extramacular Retina from Postmortem Eyes with Retinitis Pigmentosa. Investigative Ophthalmology and Visual Science, 40(1), January 1999.

[3] S. Y. Kim, S. Sadda, M. S. Humayun, E. De Juan, B. M. Melia, and W. R. Green. MORPHOMETRIC ANALYSIS OF THE MACULA IN EYES WITH GEOGRAPHIC ATROPHY DUE TO AGE-RELATED MACULAR DEGENERATION:. Retina, 22(4):464–470, August 2002.

[4] F. G. Holz, Y. Le Mer, M. M. Muqit, L.-O. Hattenbach, A. Cusumano, S. Grisanti, L. Kodjikian, M. A. Pileri, F. Matonti, E. Souied, B. V. Stanzel, P. Szurman, M. Weber, K. U. Bartz-Schmidt, N. Eter, M. N. Delyfer, J. F. Girmens, K. A. Van Overdam, A. Wolf, R. Hornig, M. Corazzol, F. Brodie, L. Olmos De Koo, D. Palanker, and J.-A. Sahel. Subretinal Photovoltaic Implant to Restore Vision in Geographic Atrophy Due to AMD. New England Journal of Medicine, page NEJMoa2501396, October 2025.

[5] T. E. Ogden and H. Ito. Avian retina. II. An evaluation of retinal electrical anisotropy. Journal of Neurophysiology, 34(3):367–373, May 1971.

[6] H. Kasi, R. Meissner, A. Babalian, H. V. Lintel, A. Bertsch, and P. Renaud. Direct localised measurement of electrical resistivity profile in rat and embryonic chick retinas using a microprobe. Journal of Electrical Bioimpedance, 1(1):84–92, December 2010.

[7] B. Wang and J. D. Weiland. Resistivity profiles of wild-type, rd1, and rd10 mouse retina. In 2015 37th Annual International Conference of the IEEE Engineering in Medicine and Biology Society (EMBC), pages 1650–1653, Milan, Italy, August 2015. IEEE.

[8] M. Jung, A. Willuweit, and V. Rincón Montes. Intraretinal Electrophysiology and Resistivity Profiles of WT and RCS Rat Retina. Sensors, 25(12):3765, June 2025.

[9] C. J. Karwoski, D. A. Frambach, and L. M. Proenza. Laminar profile of resistivity in frog retina. Journal of Neurophysiology, 54(6):1607–1619, December 1985.

[10] Sarthak Shah, Anna Kochnev Goldstein, Zhijie Charles Chen, Praful Vasireddy, Mohajeet Balveer Bhuckory, and Daniel V Palanker. Mapping the retinal resistivity with electrical impedance tomography for modeling of retinal stimulation. In Investigative Ophthalmology & Visual Science, volume 65, June 2024.

[11] S. Shah, A. Hines, D. Zhou, R. J. Greenberg, M. S. Humayun, and J. D. Weiland. Electrical properties of retinal–electrode interface. Journal of Neural Engineering, 4(1):S24–S29, March 2007.

[12] S. Mansouri, Y. Alharbi, F. Haddad, S. Chabcoub, A. Alshrouf, and A. A. Abd-Elghany. Electrical Impedance tomography – recent applications and developments. Journal of Electrical Bioimpedance, 12(1):50–62, January 2021.

[13] L. Gourdol, R. Clément, J. Juilleret, L. Pfister, and C. Hissler. Large-scale ERT surveys for investigating shallow regolithproperties and architecture, December 2018.

[14] A. Litke, N. Bezayiff, E. Chichilnisky, W. Cunningham, W. Dabrowski, A. Grillo, M. Grivich, P. Grybos, P. Hottowy, S. Kachiguine, R. Kalmar, K. Mathieson, D. Petrusca, M. Rahman, and A. Sher. What does the eye tell the brain?: Development of a system for the large-scale recording of retinal output activity. IEEE Transactions on Nuclear Science, 51(4):1434–1440, August 2004.

[15] P. Hottowy, A. Skoczeń, D. E. Gunning, S. Kachiguine, K. Mathieson, A. Sher, P. Wiącek, A. M. Litke, and W. Dąbrowski. Properties and application of a multichannel integrated circuit for low-artifact, patterned electrical stimulation of neural tissue. Journal of Neural Engineering, 9(6):066005, December 2012.

[16] P. Werginz and F. Rattay. The impact of calcium current reversal on neurotransmitter release in the electrically stimulated retina. Journal of Neural Engineering, 13(4):046013, August 2016.

[17] B.-Y. Wang, Z. C. Chen, M. Bhuckory, T. Huang, A. Shin, V. Zuckerman, E. Ho, E. Rosenfeld, L. Galambos, T. Kamins, K. Mathieson, and D. Palanker. Electronic photoreceptors enable prosthetic visual acuity matching the natural resolution in rats. Nature Communications, 13(1):6627, November 2022.

[18] A. Shin, N. Jensen, E. Butt, J. An, D. Pham-Howard, L. Galambos, K. Mathieson, T. Kamins, and D. Palanker. Amorphous silicon resistors enable smaller pixels in photovoltaic retinal prosthesis. Journal of Neural Engineering, 22(5):056017, October 2025.

