## Supplementary Information for "Retinal Resistivity Mapped by Electrical Resistance Tomography"

### Supplementary Information for:
### Retinal Resistivity Mapped by Electrical Resistance Tomography

Preprint, compiled October 13, 2026

**Anna Kochnev Goldstein[a,1], Mohammad Asif Zaman[b], Michael Aaron Sommeling[a], Sarthak Vinay Shah[c,d], Amrith Lotlikar[a], Mohajeet B. Bhuckory[b,e], Davis Pham-Howard[b,e], Zhijie Charles Chen[a], Praful Vasireddy[a], A.J. Phillips[a], Aviv Sharon[f], E.J. Chichilnisky[f], and Daniel Palanker[b,e]**

[a]Department of Electrical Engineering, Stanford University, Stanford, CA
[b]Hansen Experimental Physics Laboratory, Stanford University, Stanford, CA
[c]School of Medicine, Stanford University, Stanford, CA
[d]Department of Ophthalmology, Duke Eye Center, Duke University School of Medicine, Durham, NC
[e]Department of Ophthalmology, Stanford University, Stanford, CA
[f]Department of Neurosurgery, Stanford University, Stanford, CA

#### Literature Review

The values reported in previous studies vary by nearly two orders of magnitude, ranging from approximately 59 to 8000 Ω·cm (Table S1). One study [1] reported the resistivity for specific retinal layers, several others [2], [3], [4] as a depth-resolved graph, while others [5, 1] – as a single average value.

Table S1: A summary of the retinal resistivity values in Ω·cm reported in the literature. NFL: Nerve Fiber Layer, GCL: Ganglion Cell Layer, IPL: Inner Plexiform Layer, INL: Inner Nuclear Layer, ONL: Outer Nuclear Layer, IS and OS: Inner and Outer Segments of the Photoreceptors. This data is a subset of the previously measured values and should not be treated as a comprehensive literature analysis.

| Study | NFL+GCL | IPL | INL+ONL+IS+OS | Model |
|---|---|---|---|---|
| Karwoski et al. 1985 [1] | 7900 | 1750 | 6800 | Frog |
| Kasi et al. 2010 [2] | | 100-400 | | Rat |
| Wang et al. 2015 [3] | | 60-370 | | Mouse |
| Jung et al. 2025 [4] | | 59-289 | | Rat |
| Ogden et al. 1971 [5] | | 1200 | | Pigeon |
| Karwoski et al. 1985 [1] | | 5050 | | Frog |

#### Data Processing and Calibration

Voltage waveforms in response to injected biphasic charge-balanced pulses of current (0.5 ms per phase, anodic first) were recorded for current amplitudes ranging from 0.5-58 nA. Figure S1 shows examples of the voltage waveforms recorded in saline diluted to 1500 Ω·cm at a few sample current levels on two recording electrodes - one 30 $\mu$m away from the source (Figure S1A), and the other - at 330 $\mu$m (Figure S1B). As expected in a homogenous medium, the voltage decreases with distance L approximately as 1/L, and 330 $\mu$m away it is about 10-fold lower than 30 $\mu$m away.

The peak voltage during the anodic phase was extracted from the raw waveforms for every current level and every pair of injecting and recording electrodes. These voltages were converted to resistance by dividing them by the corresponding current and extracting the slope as R=V/I. The amplitude of the recorded voltages increased linearly with the injected current until it started reaching the saturation of the amplifiers around 1.6 mV, thus only the linear range was used to define the slope (Figure S1C). Voltages recorded at farther electrodes are lower and reach saturation at higher currents, so different current ranges were analyzed for different electrodes. For example, the voltages on the electrode in Figure S1D do not reach saturation in the measured range.

Each extracted resistance value was inherently associated with distance from the injecting electrode to the recording electrode on the MEA. As the recording electrodes were located at different distances, each injecting electrode yielded a series of resistance-distance data points. Resistance values recorded at identical distances were averaged.

To calibrate the measurement system, data was first acquired in a series of diluted saline solutions of known resistivities: 60, 500, 1500, 3000, and 5000 Ω·cm, verified with a conductivity probe (Cat. No. 89094-958; VWR International, Radnor, PA, USA). This characterization was done before and after the retinal measurement (excluding one sample where it was done only before) to account for the fact that mounting and dismounting the tissue could remove some of the brittle platinum black electroplated on the electrodes, which would affect their size and capacitance. Figure S1 shows the data for a saline dilution of 1500 Ω·cm. The same procedure was applied to all calibration dilutions and retinal samples.

##### Calibration in Diluted Saline Solutions

For each of the calibration solutions, all the resistance-distance values extracted from a single injecting electrode were plotted in the log-log scale. The data were then fitted with a -1 slope since the voltage in a homogenous conductive medium is expected to decay as 1/distance (Figure S2). The expected analytical solution is also shown in the same Figure. The difference between the measured and analytical lines reveals that injected current is lower than the nominal value, and the difference increases with the medium resistivity (Figure S2A vs. Figure S2B).

A significant fraction of current is lost due to shunting inside the MEA, and as the resistivity of the medium increases, a larger fraction of the current flows to return through this parallel shunt resistor instead of going into the medium. To ensure that the injected current can be corrected for each retinal sample of unknown resistivity, the ratio between the experimental fit

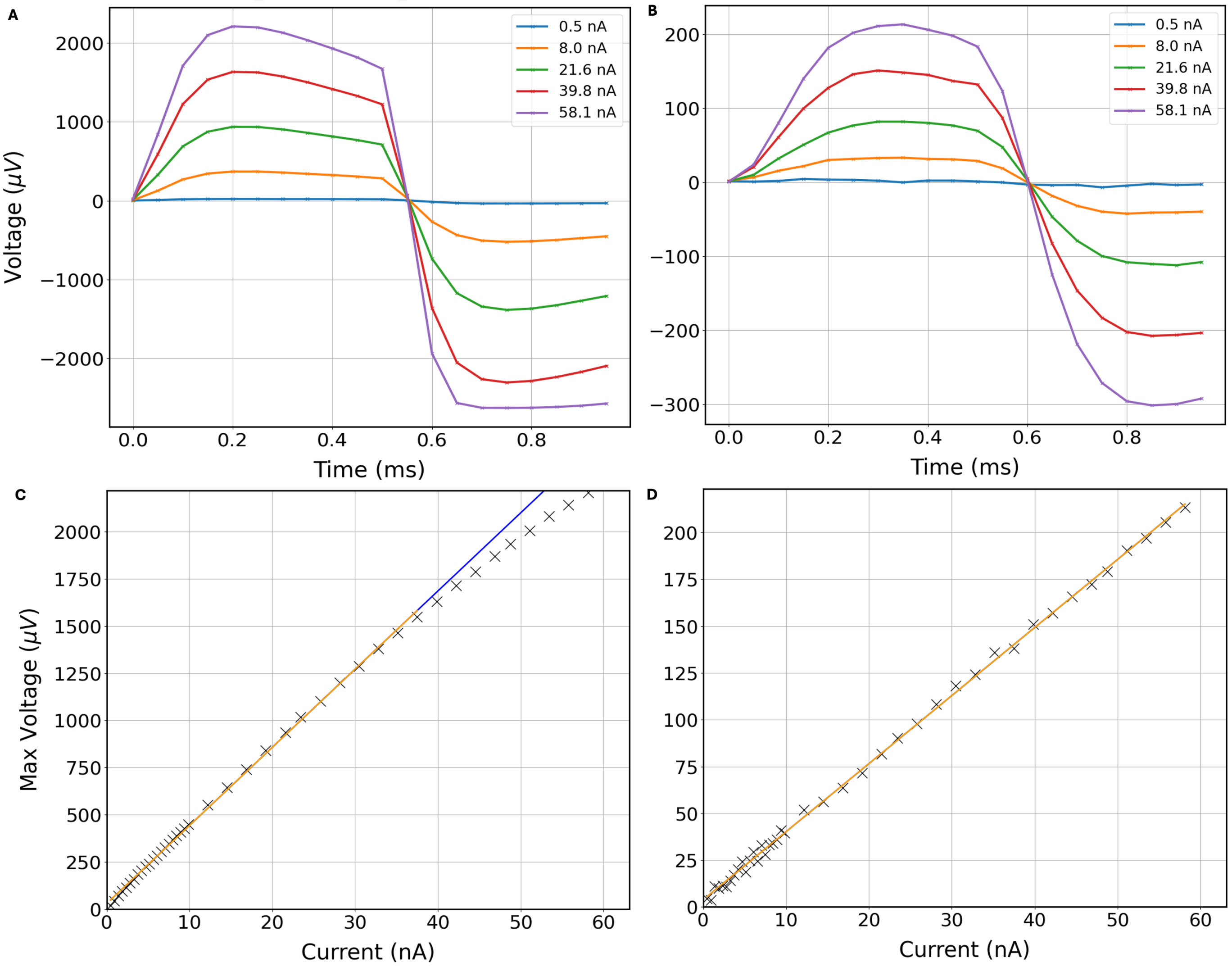


Figure S1: ***From raw voltages to resistance-distance values. A)*** *Voltage waveforms recorded in a diluted saline solution of 1538 Ω·cm at current levels ranging from 0.5 to 58 nA 30 μm away from the current source.* ***B)*** *Voltage waveforms recorded in a diluted saline solution of 1538 Ω·cm 330 μm away from the current source.* ***C)*** *Resistance between the injecting and recording electrodes extracted from the linear slope max(V)/I below saturation. For a recording electrode 30 μm away from the current source, the extracted resistance is 41.09 k*Ω*.* ***D)*** *Resistance between the injecting and recording electrodes extracted from the linear slope max(V)/I below saturation. For a recording electrode 330 μm away from the source, the extracted resistance is 3.64 k*Ω*.*

and the analytical solution was extracted across all five saline dilutions. The corresponding ratio values were then used to form a ratio-resistivity dependence curve (Figure S2C) for every injecting electrode.

The amount of current lost via the parallel shunt is defined by the ratio between the resistance of the medium and of the shunt. To monitor the resistance the injecting electrode was facing, we also recorded the potential on the injecting electrode itself (auto-potential). Divided by the injected current, it represents equivalent local resistance, as shown in Figure S2D. The dependence of the auto-resistance on the medium's resistivity combined with the current ratio dependence on the medium's resistivity established the fraction of current injected into the medium, as illustrated in Figure S3.

The deviations of the experimental data points from the linear fit in Figure S2A and Figure S2B represent the variability in the amplifier gain of the recording electrodes. As this value varied to some extent with the amplitude of the recorded signal, we refer to it as a correction coefficient rather than a fixed gain. This correction coefficient (the multiplier needed to bring the experimental data point to the linear fit) was extracted for each injecting-recording pair and each dilution, forming a correction coefficient-resistivity map.

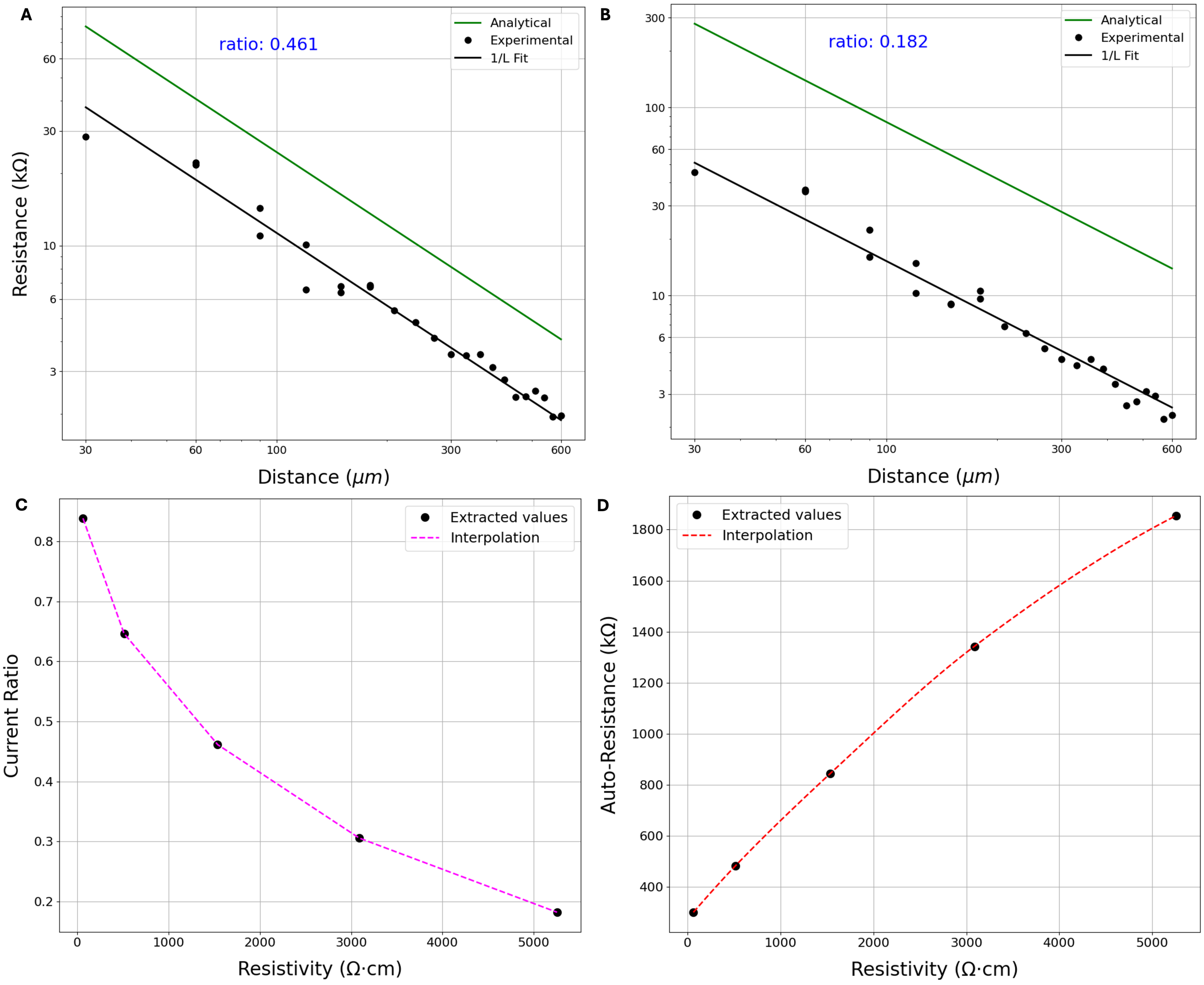


Figure S2: ***Processing calibration data. A)*** *Resistance as a function of distance from the injecting electrode (black) compared to analytical plot (green) in a solution of 1538 Ω·cm exhibits a ratio of 0.46.* ***B)*** *Resistance as a function of distance from the injecting electrode (black) compared to analytical curve (green) in a solution of 5255 Ω·cm exhibits a ratio of 0.18.* ***C)*** *Current ratio as a function of resistivity (piece-wise linear fit).* ***D)*** *Auto-resistance as a function of resistivity (monotonic fit).*

#### Retinal Samples

On retinal recordings, the calibration was performed as follows: the auto-resistance of the injecting electrode defined the equivalent resistivity of the medium sensed by the injecting electrode (Figure S3A), and the identified equivalent resistivity defined the ratio of injected current (Figure S3B). The same resistivity was also used to identify the corresponding correction coefficients for every recording electrode paired with this injecting electrode. This was done by a linear interpolation between the two closest measured resistivities (Figure S3C). After calibration, resistance values from identical distances were averaged and the retinal data was plotted as a function of distance to produce a single resistance-distance trace (Figure S4).

Each measurement line was calibrated using the described procedure twice - once using the calibration data measured before the retinal sample and once using the data measured after. As there were three measurement lines and two calibration sets, the procedure yielded up to six traces per retinal sample. All the traces from all the measurement lines and all samples of a certain rat type and orientation were averaged together to obtain the average trace and standard deviations as shown in Figure 3.

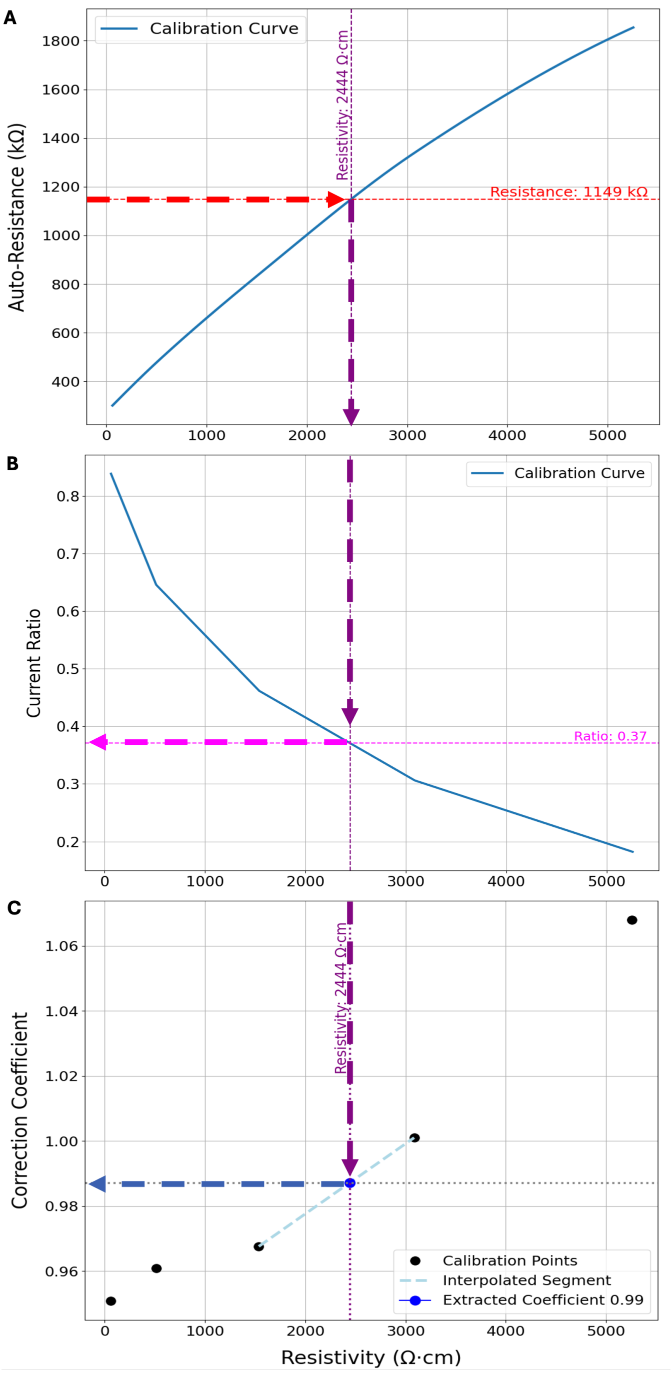


Figure S3: ***Calibrating the retinal sample.*** ***A)*** *Measured auto-resistance of 1149.1 kΩ. corresponds to an equivalent uniform resistivity of 2443.5 Ω·cm.* ***B)*** *Equivalent resistivity of 2443.5 Ω·cm corresponds to a current ratio of 0.37.* ***C)*** *Calibrating the gain coefficient of a sample recording electrode paired with the same injecting electrode as the previous panels.*

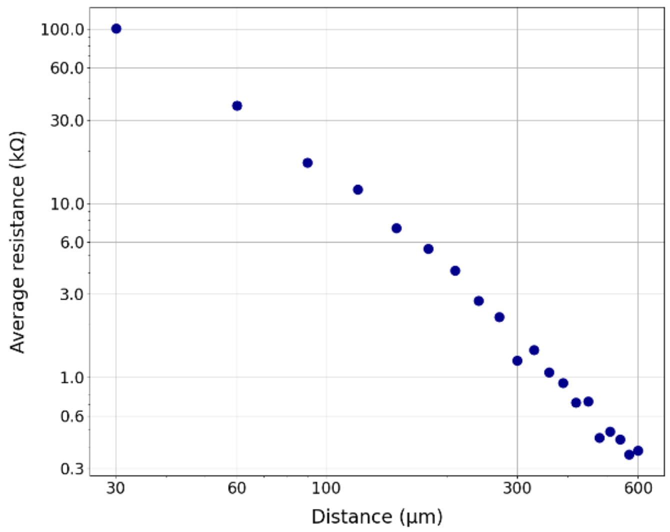


Figure S4: ***Resistance-distance trace of a retinal sample.*** *Calibrated resistance-distance trace from a single retinal sample over a single measurement line on MEA.*

#### Layer Sensitivity Analysis

To better understand how changes in the underlying resistivity profile affect the resistance-distance measurement, a sensitivity analysis was performed using COMSOL forward-modeling. First, the layers in the four-layer model were set to the resistivity values obtained by the ML decoder for the average experimental LE GCL-down trace from (3027, 1712, 2168, 939 Ω·cm), defined as the base values. Next, the resistivity of each layer was modified separately by changing it by ± 500 Ω·cm from its base value, while keeping the other layers fixed. The same simulation of injecting current from the source electrode, measuring the voltage on the recording electrodes, and extracting a resistance-distance trace was executed with all the above resistivity permutations. The resulting resistance-distance traces were plotted together using designated colors for each layer. The analysis was repeated for the LE PR-down model using the base values decoded from the GCL-down trace.

The resulting plots in Figure S5 visualize how the variation in resistivity of each layer translates into deviations of the resistance-distance trace at different distances from the current injecting electrode. The part of the trace that changed due to a perturbation in each layer is colored in the layer's designated color. For improved visibility, the sign of the applied perturbation is alternated between the layers (plus 500 Ω·cm in the NFL+GCL layer, minus 500 Ω·cm in the IPL, etc.). Figure S5B and Figure S5C show the absolute value of the same output normalized by the base resistance-distance trace, for the LE GCL-down and LE PR-down samples, respectively.

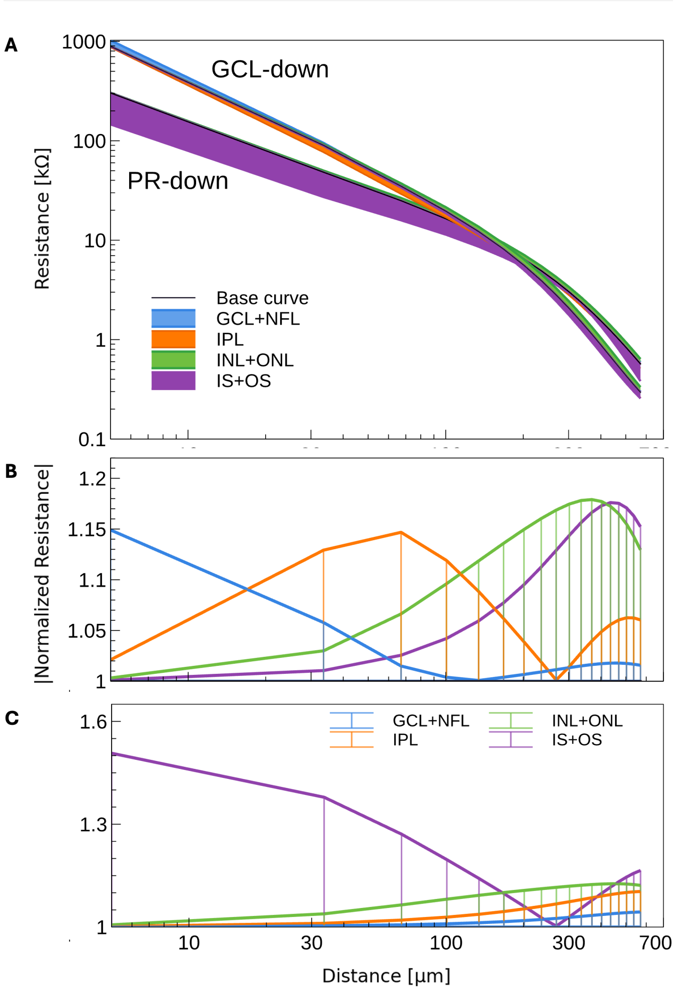


Figure S5: ***Layer sensitivity.*** ***A)*** *Color-coded resistance-distance traces for resistivity perturbations by 500 Ω·cm applied to GCL-down and the PR-down models. The plus and minus perturbations alternated between the layers for improved visibility.* ***B)*** *The same color-coded traces normalized by the base trace for the GCL-down model.* ***C)*** *The same color-coded traces normalized by the base trace for the PR-down model.*

While each change affects several regions in the resistance-distance curve, a certain part of the curve responds stronger to one layer than others. In the GCL-down orientation, each of the 4 layers has a certain dominance region, where its resistivity affects the measurement the most. At shorter distances from the injection electrode, the proximal layers dominate, and sensitivity decreases as we go deeper into the retina with larger distances. The NFL+GCL layer clearly dominates close to the injection site, up to approximately 18 $\mu$m. Since the minimal pitch of our array is 30 $\mu$m, the sensitivity of the NFL+GCL is also affected by the IPL layer. This could potentially be improved with smaller electrode pitch. The dominant range of the IS+OS layer is limited to the far-right end of the curve, and its effect significantly overlaps with the INL+OPL+ONL layer.

The PR-down orientation is dominated mainly by the IS+OS layer, most likely since this highly conductive layer in contact with MEA effectively screens the other layers. Therefore, the IPL and NFL+GCL layers, masked by the two proximal layers, are poorly detectable. Due to this limited sensitivity, the decoding of the flipped traces was not reliable. Nevertheless, the LE PR-down curve can be matched using the same values as decoded from the LE GCL-down curve. Since the NFL+GCL is more resistive than IS+OS, when the retina is placed GCL-down on the array, the measured voltage is higher than when the array faces a more conductive IS+OS layer. At larger distances, the deeper layers dominate the curve, and hence the trend reverses: resistance decreases slower with PR-down retina.

#### Resistivity Extraction from Res2DInv Traces

To convert Res2DInv's semi-continuous 1D output to a single-value resistivity for each of the retinal layers for a numerical analysis, we assumed the same 4-layer or 3-layer model for healthy and degenerated retina, respectively, with the average layer thicknesses defined from OCT imaging and histology. The 1D resistivity trace was scanned within a given depth range to identify a local extremum. If an extremum point was identified within the layer bounds, its resistivity was set as a single value for that layer. If no extrema were identified, then all the points within the boundaries were averaged. Figure S6 shows two examples of this procedure for the average experimental LE and RCS traces.

#### Accuracy Analysis

To quantify the decoders' ability to uncover the correct resistivity profiles of the 4-layer or 3-layer structures, the decoders were tested on a large synthetic data set. Using 8000 randomly generated profiles with resistivity values ranging from 60 to 6000 Ω·cm, the COMSOL forward-model resistance-distance traces were processed by both methods to decode their underlying resistivity profiles. For each trace in the test set, the decoded value of a layer was compared with its ground truth, and the average error was calculated across all the sets.

The average errors from the 4-layer traces are shown in Table S2. The ML decoder, trained on a comprehensive set of traces, yields an error increasing with depth from 1 to about 4%. Res2DInv, on the other hand, produced larger errors, also increasing with depth. In addition to loss of sensitivity with depth, Res2DInv's result is affected by the coupling between layers as the inverse problem solver tries to converge to a solution that satisfies all the given input data. Moreover, this error encapsulates the inherent inaccuracy of discretizing a continuous function into a single value for each layer.

Table S2: The average decoding error for a 4-layer retinal profile in saline.

| Decoding Method | NFL+GCL | IPL | INL+OPL+ONL | IS+OS |
|---|---|---|---|---|
| ML decoder (%) | 0.95 | 1.24 | 1.74 | 3.65 |
| Res2DInv (%) | 10.1 | 9.6 | 36 | 62.2 |

Running a similar accuracy test on the 3-layer structures generally follows a similar trend (Table S3), albeit the accuracy starts dropping much sooner. This reflects the ML decoder's

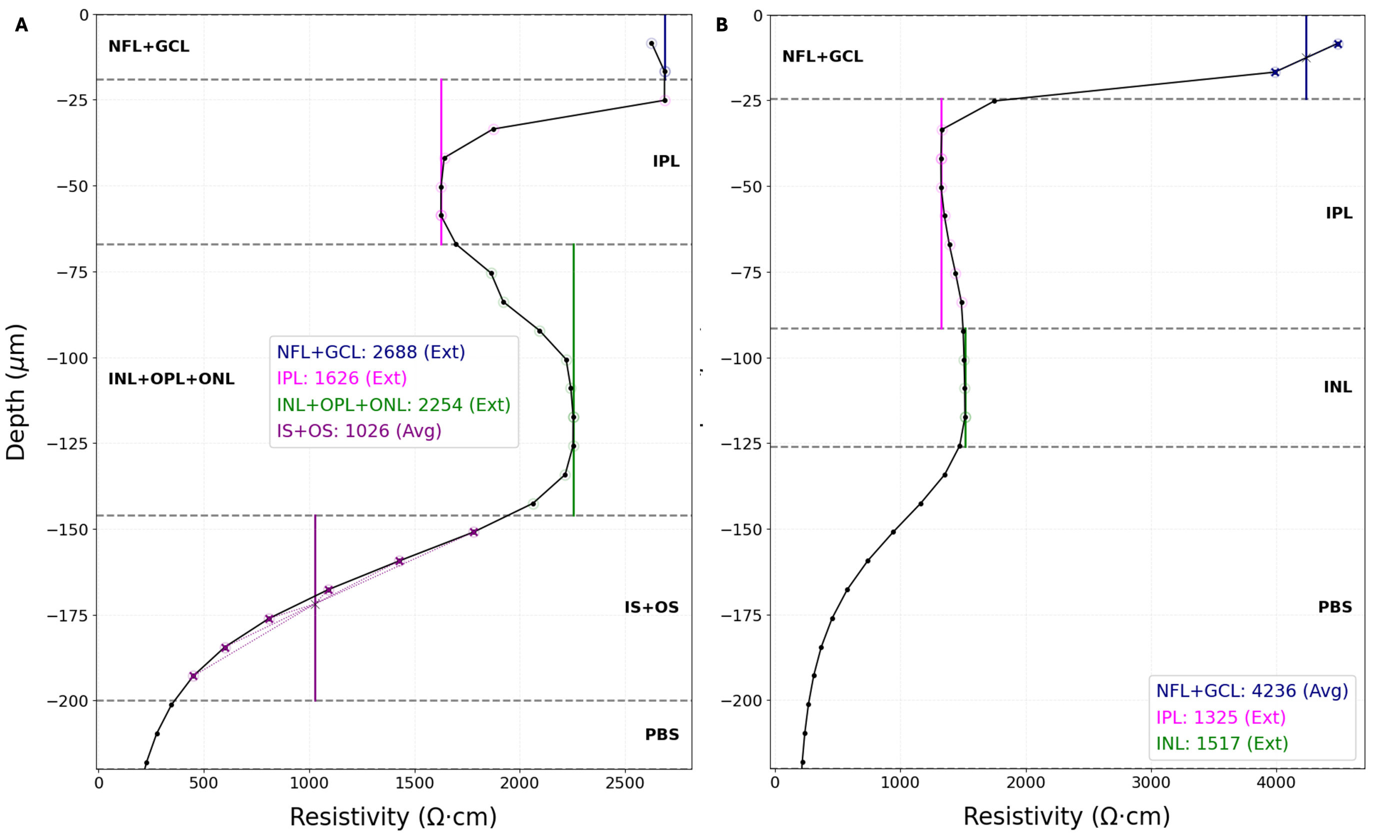


Figure S6: ***Resistivity extraction from Res2DInv's output trace.*** *Right: The decoded output and extracted values for the average LE trace. Left: The decoded output and extracted values for the average RCS trace.*

reduced accuracy given fewer tuning parameters, as well as Res2DInv's limited ability to accurately decode profiles with high contrast between adjacent layers, especially when a thinner INL interfaces a very conductive saline layer.

Table S3: The average decoding error for a 3-layer retinal profile in saline.

| Decoding Method | NFL+GCL | IPL | INL |
|---|---|---|---|
| ML decoder (%) | 1.67 | 3.24 | 8.37 |
| Res2DInv (%) | 10.5 | 25.8 | 56 |

#### References

[1] C. J. Karwoski, D. A. Frambach, and L. M. Proenza. Laminar profile of resistivity in frog retina. *Journal of Neurophysiology*, 54 (6):1607–1619, December 1985.

[2] H. Kasi, R. Meissner, A. Babalian, H. V. Lintel, A. Bertsch, and P. Renaud. Direct localised measurement of electrical resistivity profile in rat and embryonic chick retinas using a microprobe. *Journal of Electrical Bioimpedance*, 1(1):84–92, December 2010.

[3] B. Wang and J. D. Weiland. Resistivity profiles of wild-type, rd1, and rd10 mouse retina. In *2015 37th Annual International Conference of the IEEE Engineering in Medicine and Biology Society (EMBC)*, pages 1650–1653, Milan, Italy, August 2015. IEEE.

[4] M. Jung, A. Willuweit, and V. Rincón Montes. Intraretinal Electrophysiology and Resistivity Profiles of WT and RCS Rat Retina. *Sensors*, 25(12):3765, June 2025.

[5] T. E. Ogden and H. Ito. Avian retina. II. An evaluation of retinal electrical anisotropy. *Journal of Neurophysiology*, 34(3):367–373, May 1971.
